# Connectivity and population structure in a marginal sea – a review

**DOI:** 10.1101/2024.11.11.622907

**Authors:** Simon Henriksson, Per Erik Jorde, Charlotte Berkström, Guldborg Søvik, Pierre De Wit, Halvor Knutsen, Even Moland, Carl André, Marlene Jahnke

## Abstract

The current biodiversity crisis calls for conservation measures that limit the negative human impact on important habitats and sensitive wild populations. To effectively protect biodiversity at all levels, including intraspecific diversity, conservation measures should be aligned with the connectivity and genetic structure of wild populations. In this review, we synthesise scientific literature on the connectivity and population structure of marine species in the Skagerrak – a marginal sea in the northeast Atlantic Ocean. We discuss the results in relation to the current management practices in the region, as well as the general transferability of our findings. The Skagerrak is one of the most intensively studied regions within this research field, and our findings show that the overall connectivity with adjacent seas is high, but asymmetric, for most species. Simultaneously, most species have populations in the Skagerrak distinct from both each other, and those in adjacent seas.

Most of this population structure is associated with the convoluted Skagerrak coastline – population structure is common both among coastal populations and between coastal and offshore populations. In many mobile species, multiple populations are temporally sympatric in certain areas, but retain their genetic divergence through natal homing or other barriers to gene flow. The presence of population structure despite high connectivity is a challenge for area-based protection measures, and calls for temporally flexible management that also monitors intraspecific genetic diversity on multiple timescales.

## 1. Introduction

Biodiversity loss is an ongoing crisis, that negatively impacts both global and local ecosystems (Cardinale *et al*., 2012). The currently elevated extinction rate for wild species is often referred to as “the sixth mass extinction” (Cowie *et al*., 2022) and is caused by various anthropogenic pressures, such as climate change, habitat fragmentation, and overexploitation of wild populations (Pievani 2014; Ceballos *et al*., 2015). The loss of species is likely preceded by losses of intraspecific diversity (Ceballos *et al*., 2017), and it has consequently been argued that the population, rather than the species, is the relevant unit for conservation (Reydon 2019; Allendorff *et al*., 2022). Populations with large population sizes and high genetic diversity are more resilient to environmental changes, and they are more likely to harbour alleles that may prove beneficial in future environments – referred to as evolutionary potential (Bürger & Lynch, 1995; Frankham *et al*., 1999; Allendorf *et al*., 2022). For these reasons, the importance of conserving genetic diversity within species is recognised increasingly in international conventions and legislation, including Goal A and Target 4 in the recent Kunming-Montreal Global Biodiversity Framework (CBD/COP/15/L.25; Convention on Biological Diversity, 2022). Area-based protection, such as marine protected areas (MPAs), is highlighted as one of the main tools to prevent biodiversity losses, especially in Target 3, stating that 30 % of land and water areas should be protected by year 2030 (Convention on Biological Diversity, 2022). To be effective, however, area-based protection should build on a sound understanding of the spatial requirements of the target populations, and the connectivity among different populations (Goetze *et al*., 2020; Beger *et al*., 2022).

Connectivity refers to the exchange of energy, biomass, or genetic material between populations or geographic locations (Selkoe *et al*., 2016). There are several types of connectivity, which can be conceptualised in different ways. In this review, we use a conceptual connectivity framework consistent with, e.g., Lowe & Allendorf (2010), Gagnaire *et al*. (2015), Selkoe *et al*. (2016), and SEA- UNICORN (n.d.), depicted in Figure 1, and detailed in Text Box 1. Connectivity can promote the persistence of species and intraspecific genetic diversity through gene flow, but can also assist in restoring biodiversity at all levels after it has been depleted, through the dispersal and movement of organisms and material across populations, communities, and ecosystems (Balbar & Metaxas, 2019). A related concept is population structure, the subdivision of species into multiple populations (Waples & Gaggiotti, 2006; see also Text Box 1), a pattern which can only arise and persist when genetic connectivity between populations is low (Slatkin 1987). Differentiated populations within a species may harbour different genetic adaptations, enabling them to occupy different habitats or ecological niches (Stronen *et al*., 2022).

**Figure 1.**
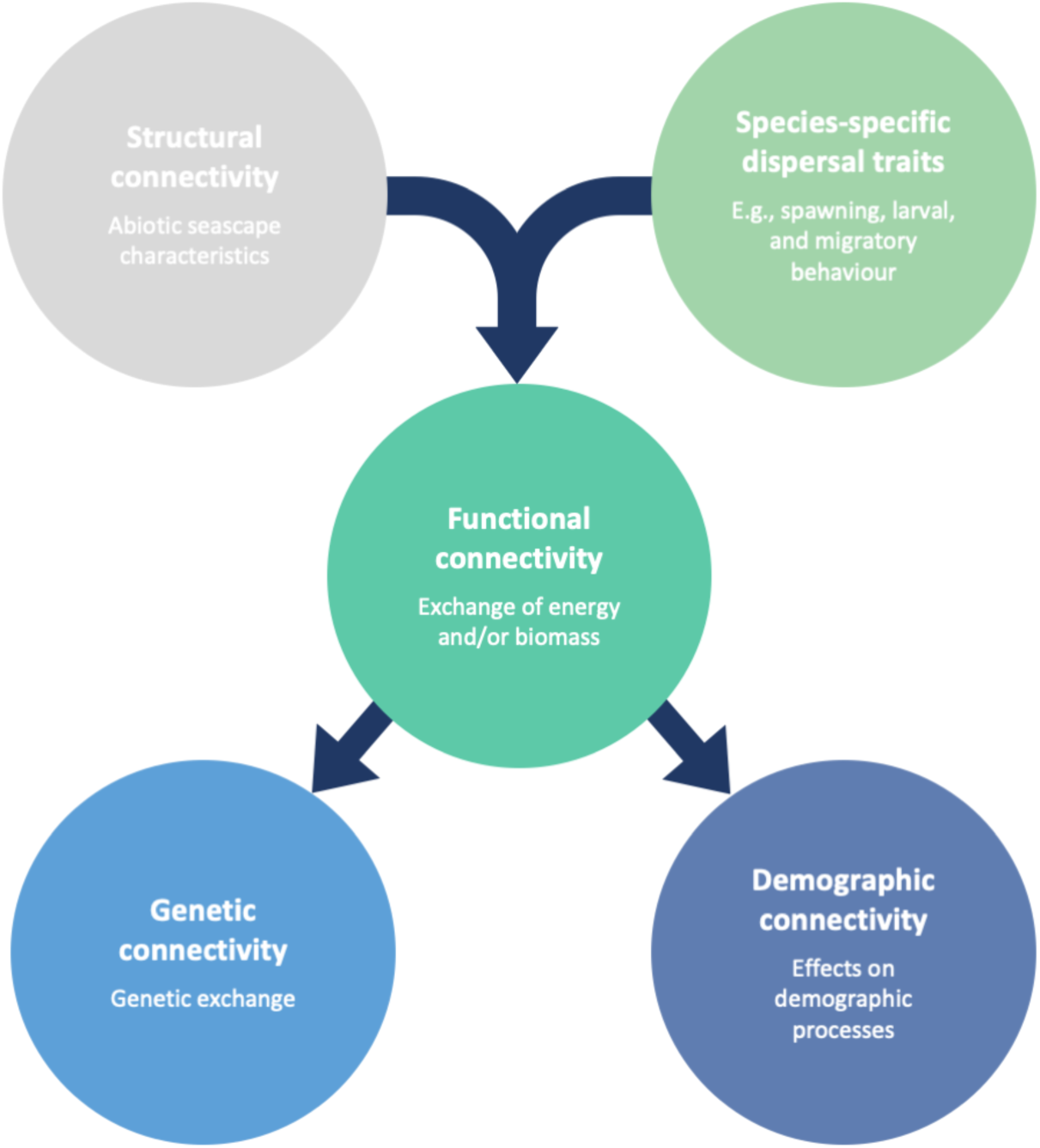
Concept map visualising the relationships of different types of connectivity. The structural connectivity of the seascape interacts with species-specific dispersal traits to shape functional connectivity patterns – actualised exchange of biomass or energy between populations or locations. Functional connectivity does not necessarily lead to genetic connectivity (gene flow) or demographic connectivity (effects on population-level processes) between populations, but is a prerequisite for both.

In addition to their evolutionary significance, connectivity and population structure also have direct applications in conservation and stock management (Riginos & Beger 2022; Jahnke & Jonsson 2022). For instance, knowledge on both the connectivity and population structure of managed species is essential to accurate delineation of management units (e.g., fish stocks, see Reiss *et al*., 2009), as the distribution ranges of marine populations can vary widely, from trans-oceanic to highly local (Kerr *et al*., 2017), as well as overlap (e.g., Aarestrup *et al*., 2022). Still to this day, management units are often geographically delineated based on legislative borders between nations or regions, when they should ideally be based on species biology and delineate demographically and/or genetically independent populations (Palsbøll *et al*., 2007). Similarly, area-based protection measures such as marine protected areas (MPAs) should, ideally, cover a sufficient number of important, and well-connected, habitat patches to ensure the persistence of populations (Goetze *et al*., 2020; Beger *et al*., 2022). For instance, prioritising the protection of areas with both high total larval contribution to adjacent sites, and high self-recruitment, has been shown to be effective in fisheries management (Krueck *et al*., 2022).

Making these prioritisations, however, requires sufficient knowledge on the connectivity of populations in the area, to elucidate the meta-population structures and source-sink dynamics of populations (Beger *et al*., 2022). Synthesising information on both population structure and connectivity is, thus, crucial in establishing a biological baseline on which informed management actions can be based.

This literature review aims to summarise biological knowledge on connectivity and population structure in marine species in the Skagerrak – a marginal sea in the northeast Atlantic. The results are of fundamental relevance for both local and regional management, especially in assigning management units, and designing MPA networks. Due to the extensive research performed in this area, we also discuss how conclusions from the Skagerrak may be transferable to other systems, potentially enabling more general conclusions about the relationship between population structure and connectivity in marine species.

### Text Box 1: Connectivity and population structure

#### Connectivity

Fundamentally, there is **structural connectivity** – the connectivity of the seascape itself, which is the outcome of seascape features such as bathymetry, the shape of the coastline, and ocean currents at various depths. How this structural connectivity affects the connectivity of species depends on the **biological dispersal traits**, specific to the species or populations studied. The interactions of structural connectivity and biological dispersal traits creates **functional connectivity**, i.e., the realised exchange of biomass between populations or geographic locations, most commonly referring to the dispersal of individuals. However, dispersed individuals may not be considered as “recruited” to the new populations if they do not remain in the new population until reaching a certain age or size, or contribute with genetic material. Thus, functional connectivity is distinct from both **demographic connectivity** and **genetic connectivity**. **Demographic connectivity** describes the extent to which immigration and emigration affect demographic processes, such as population growth, and **genetic connectivity** refers to gene flow between populations, affecting evolution and adaptation (Selkoe *et al*., 2016). Importantly, connectivity may be asymmetric, meaning the flow of energy, biomass, and/or genetic material can be higher in one spatial direction than the other (Beger *et al*., 2010). This is particularly true in marine environments, where ocean currents often have one prevailing direction.

#### Connectivity barriers

Areas where connectivity is distinctly lower than in surrounding regions are commonly referred to as connectivity breaks or **connectivity barriers**. Depending on the type of connectivity assessed, such barriers may be caused by either seascape features (structural connectivity; Selkoe *et al*., 2016), strong environmental gradients (Johannesson & André, 2006), prezygotic isolation due to behavioural differences – e.g., assortative mating (Schumer *et al*., 2017) and natal homing (André *et al*., 2016) – or postzygotic incompatibilities (Orr & Turelli, 2001).

#### Population structure

**Population structure** refers to the subdivision of species into more-or-less distinct groups of individuals, or “populations” (Waples & Gaggiotti, 2006). This subdivision can vary in strength, depending on environmental factors, the species’ biology, and the geographical area. Population structure tends to be less pronounced in marine species, as they often have both large effective population sizes and high dispersal potential (e.g., Ward *et al*., 1994). While population structure is commonly referred to in population genetic terms, it can be assessed in a multitude of ways – including chemical isotope analysis, and morphometry – in addition to genetic methods.

## 2. Methods

### 2.1. Study area

The Skagerrak is a small sea sharing its borders with Norway, Denmark and Sweden. The area is characterised by the Norwegian Trench, following the Norwegian coast, with depths down to 700 m in the eastern Skagerrak and a deep sill in the west towards the North Sea of approximately 270 m depth (Rohde, 1996). The Norwegian and Swedish coastlines are convoluted and topographically complex, characterised by many small inlets and fjords, as well as the long Oslofjord, which extends approximately 90 km northwards. Together with the Kattegat and Danish Straits, the area is often referred to as a “transition zone” between the near-oceanic North Sea and the brackish Baltic Sea (Gustafsson & Stigebrandt, 1996). The southern Skagerrak receives an inflow of water from the North Sea, turning in a counter-clockwise direction along the Swedish and Norwegian Skagerrak coasts (Danielssen *et al*., 1997). Brackish water from the Baltic Sea flowing northward along the Swedish west coast also enters the Skagerrak and mixes with the North Sea water to form the Norwegian Coastal Current (Gustafsson & Stigebrandt, 1996). The Skagerrak has received considerable attention from population genetic and evolutionary research, due to the presence of strong environmental gradients and genetic clines in multiple marine species (Johannesson *et al*., 2020). The high structural connectivity with the North Sea and the Baltic Sea has also bred interest in connectivity research on marine species in the area.

The Skagerrak is one of the world’s most productive oceans (Olsson, 1993), but both offshore and coastal Skagerrak ecosystems have been strongly impacted by anthropogenic pressures such as bottom trawling, depletion of fish stocks, and trophic imbalances (e.g., Svedäng & Bardon, 2004; Baden *et al*., 2012; Eigaard *et al*., 2016). Despite this, both implementation and holistic assessment of MPAs in this region is considerably lacking, compared to similar marine regions (Roessger *et al*., 2022).

Synthesising the current scientific knowledge on connectivity and population structure is an essential step toward establishing effective regional management practices, conserving genetic diversity and rebuilding depleted stocks. Similar to how the neighbouring Baltic Sea has been argued to be a good case study for future climate change in the ocean (Reusch *et al*., 2018), the intensive research on connectivity and population structure in the Skagerrak makes this region an excellent case study of connectivity in a marginal sea.

### 2.2. Literature search

We performed a systematic literature search for studies on connectivity and population structure in the Skagerrak on the Web of Science database on the 3rd of May 2023. Our search string was structured in three main sections, defining a) the area of study, b) the context, and c) the methodological approach. The search string in full was:

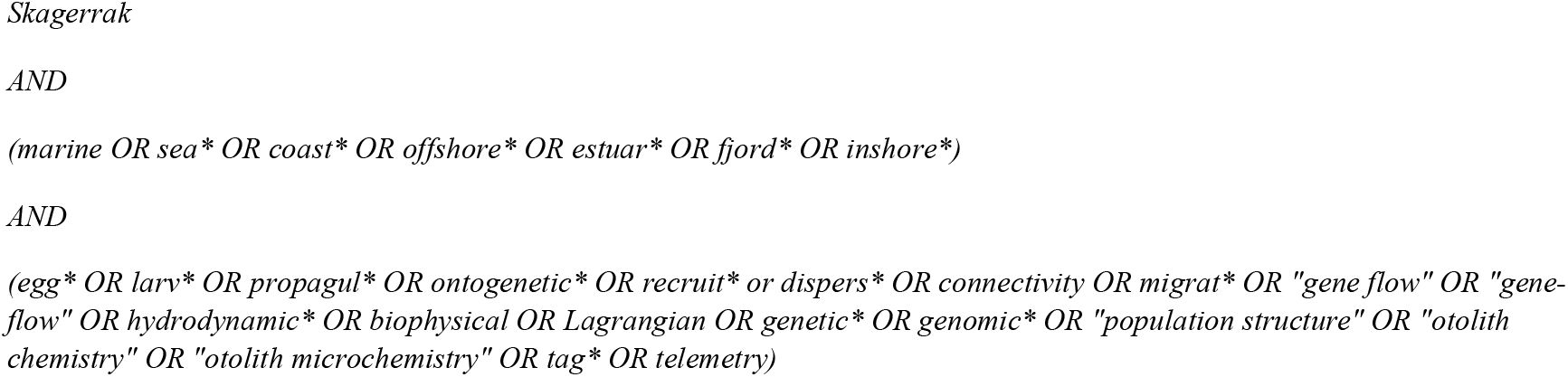

The length of the third section reflects the highly variable methodological approaches available to assess connectivity and population structure. The Web of Science database only searches through the Title, Keywords, and Abstract of the original publications. Hence, we supplemented the list of publications from the systematic literature search by manually adding scientific publications of relevance to this review.

### 2.3. Screening

The full list of publications was screened according to a set of five exclusion criteria (A-E; see Table 1). Publications were excluded if they A) had a non-marine context; B) were not in the Skagerrak; C) did not investigate connectivity of any marine species; D) were a review, meta-analysis, short-format, or non-peer-reviewed article; or E) were inaccessible.

**Table 1.**
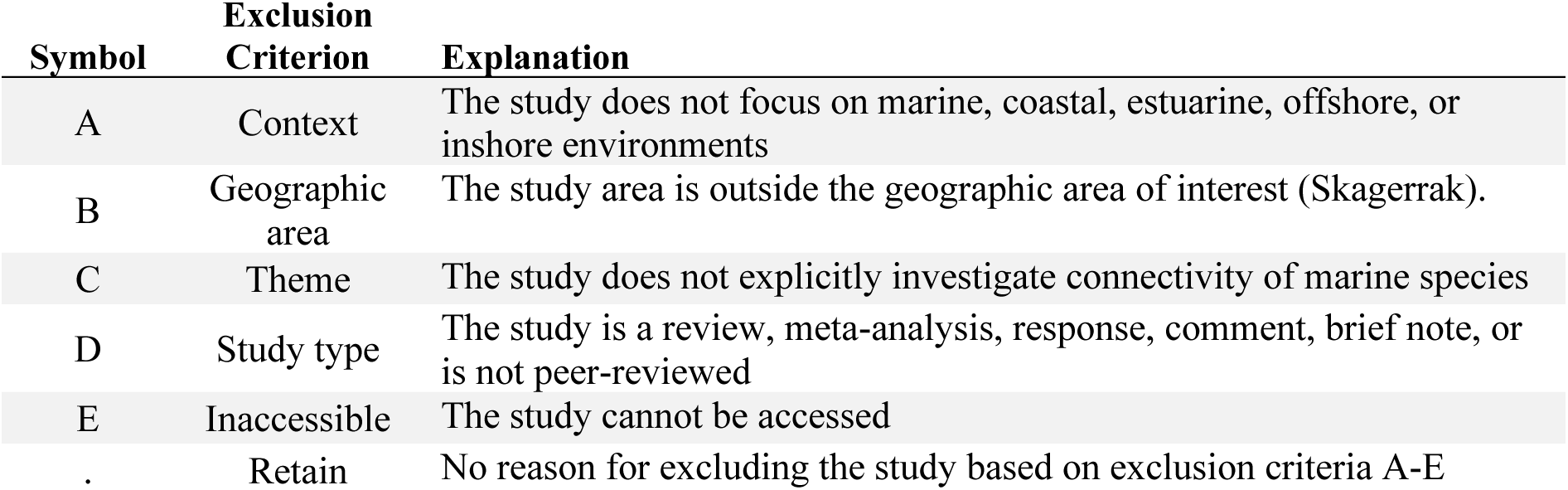
Exclusion criteria used for the screening of publications.

The systematic literature search yielded a list of 413 unique scientific publications. Out of these, 113 (27 %) were eligible for review. Most excluded publications were so based on thematic irrelevance, i.e., not explicitly assessing connectivity in marine species (exclusion criterion C; 58 %). We supplemented the list of 113 publications by manually adding 59 relevant publications that the authors were aware of, or that were cited in reviewed publications. Thus, after screening, a total of 172 scientific publications were assessed as eligible for review (Figure 2A).

**Figure 2.**
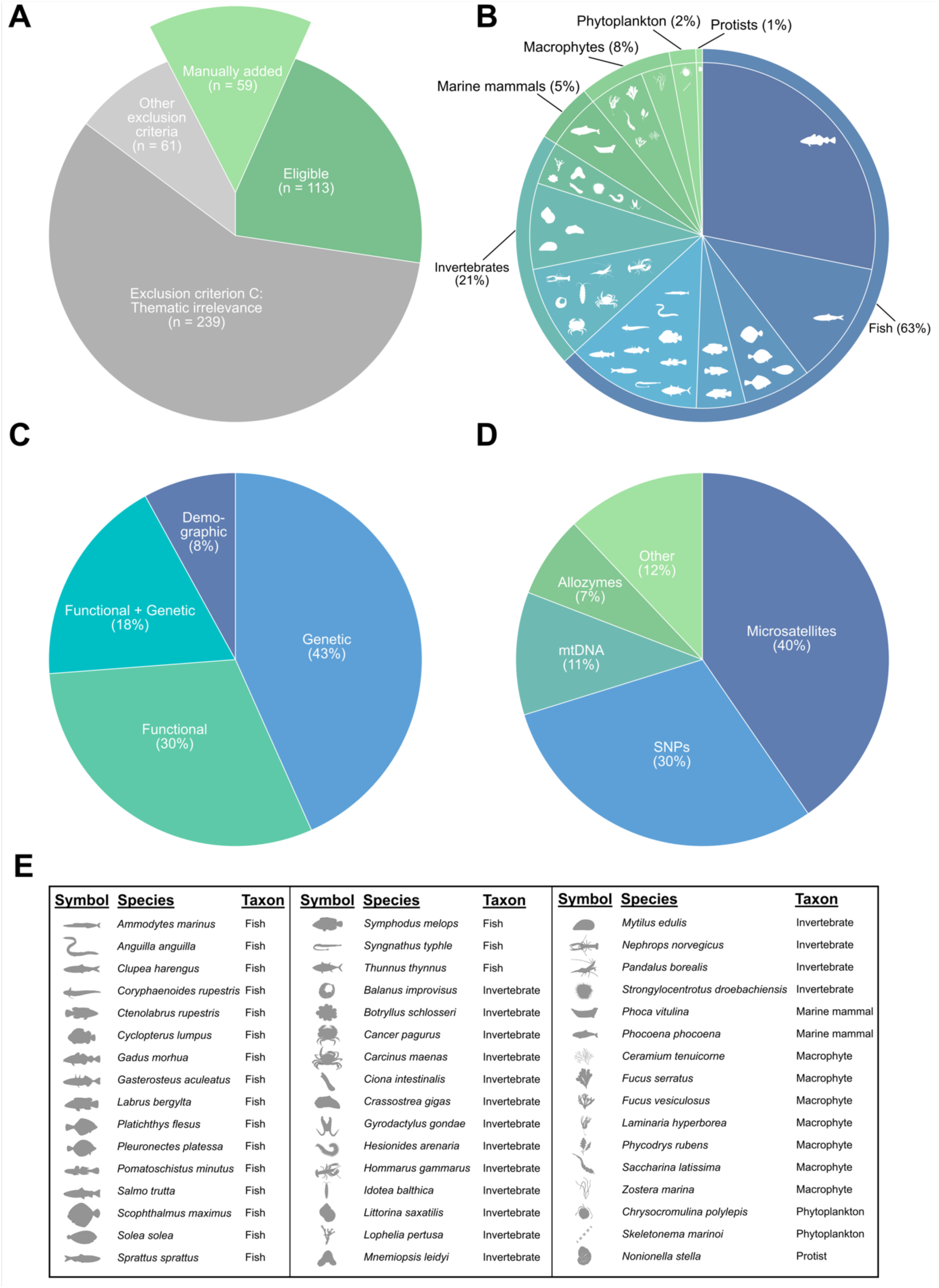
Summary of the scientific publications assessed in this review. **A)** Overview of the 413 publications from the systematic literature search on Web of Science, with pie slices representing the proportion deemed eligible (green) or ineligible (red) for this review. The 59 manually added publications are represented as the proportion in relation to the 413 publications in the systematic literature search. Subplots **B-D** summarise the studies included in this review: **B)** the relative numbers of studies per taxon (outer pie chart) and species or species group (inner pie chart); **C)** the types of connectivity assessed; and **D)** which genetic marker types have been used in the genetic studies. In cases where a single publication fit multiple categories (e.g., a publication studied two different species), the publication has been represented as multiple studies in subplots **B-D** (e.g., one study on species I and one study on species II). Subplot **E)** lists all 48 species included in this review.

### 2.4. Data extraction

Publications were divided amongst the authors, who extracted information on the study design and methodology, and summarised the relevant results. We sorted publications into three categories, based on both their methodology and the types of results presented. The categories were: publications assessing a) contemporary connectivity, b) geographically defined barriers to connectivity, and c) population structure. Note that these categories are not mutually exclusive – a single publication may assess all three, in concert.

In the *contemporary connectivity* category, we included publications assessing either active (migration) or passive dispersal (e.g., egg and larval drift) within a single generation. We included publications assessing connectivity both to, from, and within the Skagerrak. The main methodological approaches to assess contemporary connectivity were tagging studies (mark-recapture and acoustic telemetry) and biophysical modelling studies. We only included publications that provided explicit information on dispersal direction, dispersal distance, or home range in this category. The contemporary connectivity publications were further grouped into two subcategories based on the geographic scale being assessed. The first subcategory assessed connectivity on larger scales, between the Skagerrak and the North Sea, the Kattegat, and/or the Baltic Sea. The second subcategory assessed connectivity on smaller scales – here defined as limited to within the Skagerrak and smaller, i.e., mostly at scales of a few kilometres.

Publications were sorted into the *geographic connectivity barrier* category if they inferred or specifically assessed geographically defined barriers to connectivity. Inference methods varied across publications, but included both genetic methods to detect distinct genetic barriers, and biophysical modelling approaches to detect areas where biophysical barriers limit connectivity.

Publications were included in the *population structure* category if they assessed the presence of distinct groups of individuals, or variation in specific traits, within species. The methodological approaches to assess population structure differed among publications within this category, but most publications used population genetics, morphometry, or chemical isotope analyses. Publications were then further sorted into subcategories based on the number of sampled sites in the Skagerrak, and whether they described population divergence between the Skagerrak and adjacent seas (the North Sea, Kattegat, or Baltic Sea), or within the Skagerrak. Although analysing population structure also provides insight into genetic connectivity of populations, particularly over longer evolutionary timescales (Kool *et al*., 2013), in this review, we have chosen to consider population structure separately from contemporary connectivity (functional, demographic, and genetic), as these concepts have slightly different implications for conservation management.

### 2.5. Barrier analysis

We used BARRIER 2.2 (Manni *et al*., 2004) to re-analyse genetic distance estimates and infer potential genetic barriers. Specifically, we included studies on native species with at least three sampling sites in the Skagerrak, which provided both pairwise *F*ST or *G*ST estimates and geographic coordinates for the sampling sites (Supplementary Table S1). The study area, as defined in BARRIER, was between latitudes 55.6 and 60.0°N, and longitudes 4.0 and 13.2°E. This slightly exceeds the geographic limits of the Skagerrak, thus, providing some geographic context to potential barriers within the Skagerrak. Analyses were run separately for each study, and the number of barriers, which must be set *a priori* in BARRIER, ranged from 1 to 6. The number of barriers was assigned based on the number of study sites in the Skagerrak, as detailed in Supplementary Table S2. Note that this analysis did not explicitly account for mechanical admixture of multiple populations at a single sampling location, as many of the original publications did not. Similarly, statistical significance of inferred barriers could not be assessed since this information requires replicate divergence data, e.g., separate *F*ST values from different loci, which was not available for most studies.

## 3. Results

### 3.1. Literature summary

Connectivity and population structure is a research field that has been gaining interest in recent years in Skagerrak species (Figure 3). The first included study was published in 1990, and the number of studies per year have increased gradually since, especially during the 2000s and 2010s. At present, around 10 studies on connectivity and population structure are published every year.

**Figure 3.**
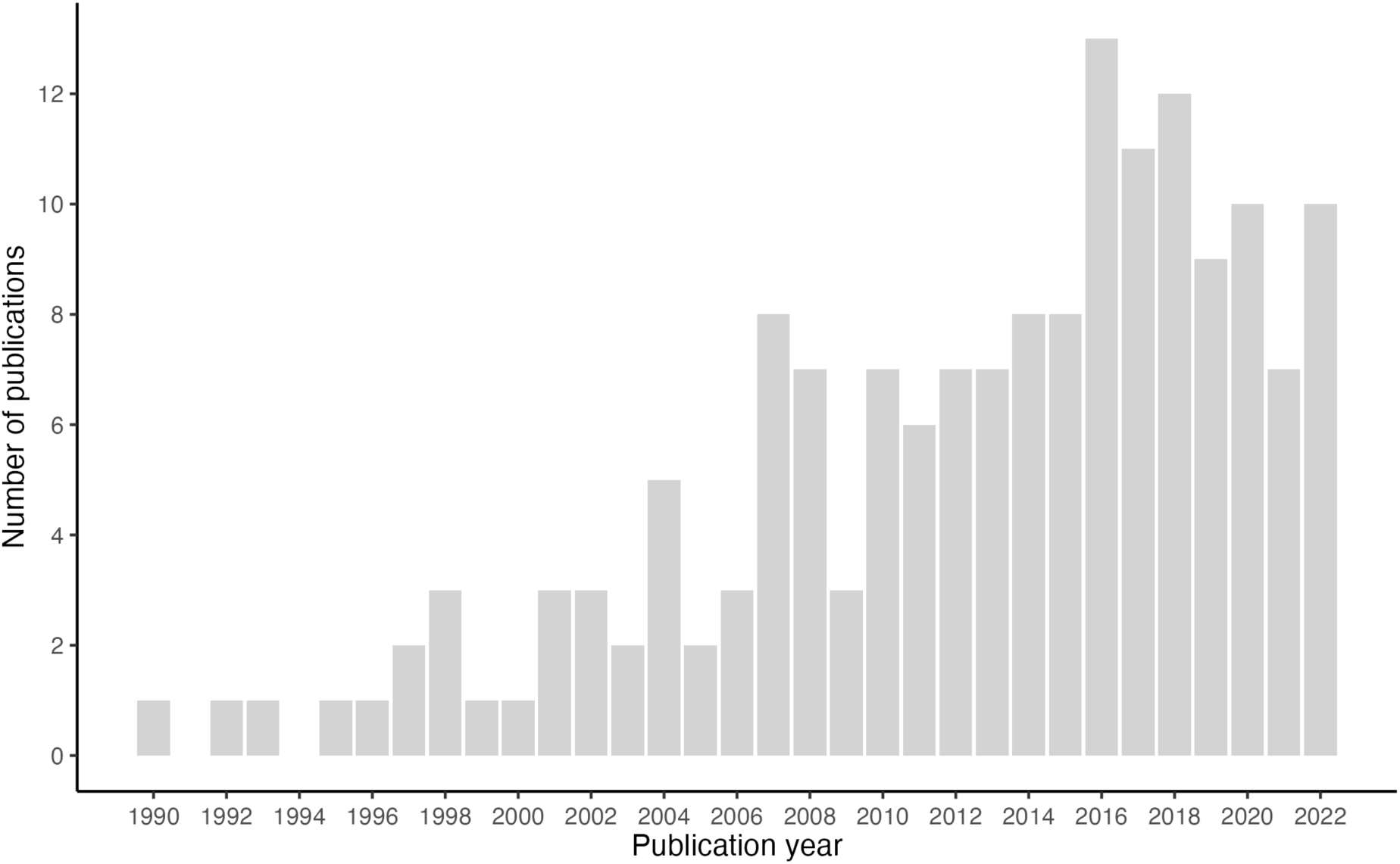
Publication years of the studies included in this review. The bar heights represent the numbers of studies, eligible for this review, published each year. As the literature search was performed in March 2023, studies published in 2023 are excluded from this figure. Hence, only results from whole years are displayed.

In total, population structure or connectivity has been assessed in 48 unique Skagerrak species (Figure 2E). The scientific literature on population structure and connectivity in Skagerrak species is strongly dominated by fish species (63 % of studies; Figure 2B). In fact, Atlantic cod (*Gadus morhua*; 28 %) and Atlantic herring (*Clupea harengus*; 11 %) are the subject of investigation in 39 % of the studies. Other fish taxa represented are flatfishes (6 %) and labrids (5 %), together with other fish species (13%). Invertebrates are the next most represented taxon (21 %), with mainly crustacean (9 %) and mollusc species (8 %) studied, but also other invertebrate species (4 %). The marine mammals, harbour porpoise (*Phocoena phocoena*; 4 %) and harbour seal (*Phoca vitulina*; 1 %), make up 5 % of studies. Macrophytes are the subject of 8 % of studies, with 5 % on macroalgae and 3 % on eelgrass (*Zostera marina*). Phytoplankton and protists are the least studied taxa in the Skagerrak, covered by 2 and 1 % of studies, respectively, and represented by three species in total (*Chrysochromulina polylepis*, *Skeletonema marinoi*, and *Nonionella stella*).

Most studies have assessed genetic (43 %) or functional connectivity (30 %), but 18 % of studies have also studied both in concert. Fewer studies (8 %) have assessed demographic connectivity (Figure 2C). Studies using genetic tools to assess population structure and connectivity differ with regard to the types of genetic markers used (Figure 2D). Overall, microsatellite (40 %) and single-nucleotide polymorphism (SNP) loci (30 %) are the most common, however, methods have changed substantially over time. The use of SNPs has grown in popularity since the mid-2010s, and has almost completely replaced other markers in recent years. Of the publications included in this review, 93 assessed contemporary connectivity (55 %), 12 assessed geographic barriers to connectivity (7 %), and 131 assessed population structure (78 %).

### 3.2. Contemporary connectivity

#### 3.2.1. Connectivity with adjacent seas

Connectivity is generally high between Skagerrak populations and populations in the North Sea, Kattegat, and Baltic Sea (Figure 4). Although contemporary connectivity has been assessed for much fewer species (n=18) than population structure has, the reviewed literature points toward high connectivity between the seas. Connectivity into the Skagerrak is high in most assessed species, both from the west (North Sea: 92 % of species) and the south (Kattegat: 100 %; and Baltic Sea: 83 % of species; Figure 4A). Connectivity out of the Skagerrak is, similarly, high into the North Sea (100 %), but slightly fewer species show connectivity in the southward direction (Kattegat: 75 %; and Baltic Sea: 67 %; Figure 4B). There are, however, fewer studies assessing southward connectivity rather than northward connectivity in the Kattegat-Baltic Sea geographic region, likely due to the prevailing ocean current patterns. The main direction of passive dispersal is in the northward direction for organisms with shallow-drifting propagules, and southward for those with deeper drift depths (Pacariz *et al*., 2014a; 2014b; Jonsson *et al*., 2016).

**Figure 4.**
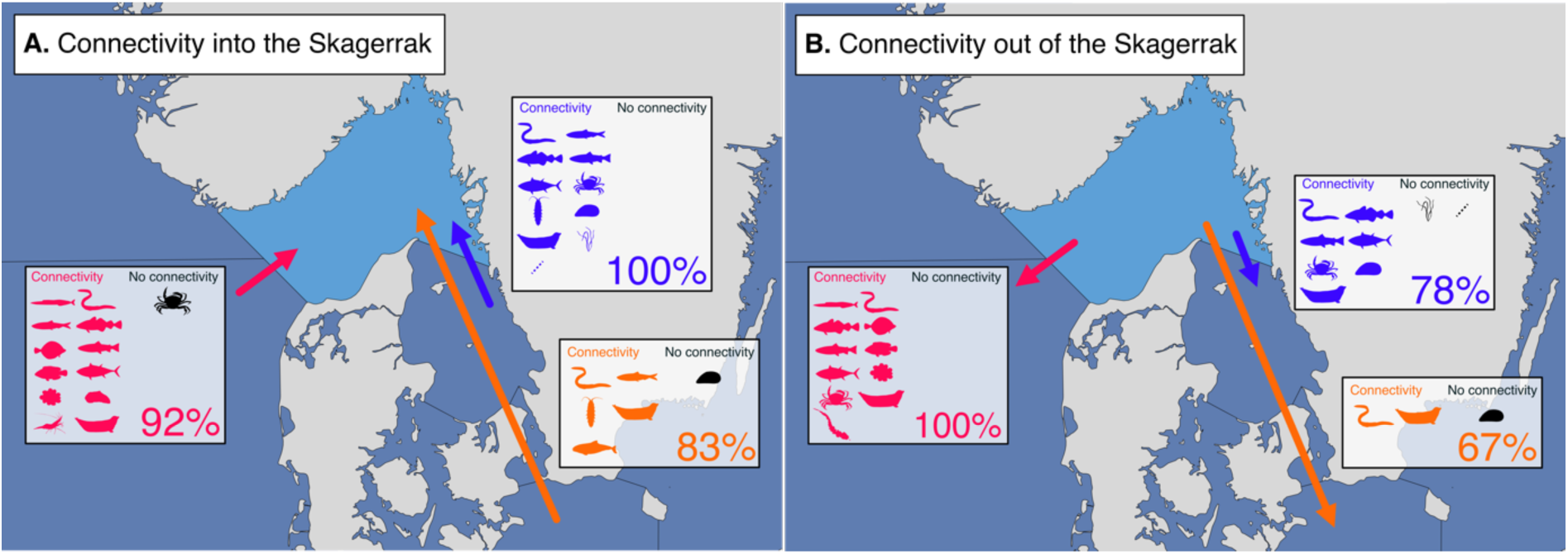
Contemporary connectivity of Skagerrak species with the adjacent North Sea (pink), Kattegat (blue), and Baltic Sea (orange). The figure summarises functional, genetic, and demographic connectivity within one generation **A)** into, and **B)** out of the Skagerrak. The boxes show which species have been assessed, and the proportion of these species for which connectivity has been found (in colour). A key for the species symbols is found in Figure 2E.

##### Species with active dispersal

For European eel (*Anguilla Anguilla*), Atlantic cod, and Atlantic herring, large amounts of eggs and larvae are passively brought into the Skagerrak from populations spawning in the North Sea or further out in the Atlantic Ocean (Stenseth *et al*., 2006; Ruzzante *et al*., 2016; Berg *et al*., 2017). Some adults later perform active migrations westwards out of the Skagerrak after reaching a certain size or age, as documented for eel and cod (Westerberg *et al*., 2014; Pihl & Ulmestrand 1993; André *et al*., 2016) and implied for herring (Ruzzante *et al*., 2006). In cod, a large proportion of eggs and larvae also passively drift into the Skagerrak from the Kattegat and Danish straits (Jonsson *et al*., 2016; Barth *et al*., 2017), and active southward migration of adult cod also occurs, but is less well-documented (André *et al*., 2016; Hüssy *et al*., 2021). For Atlantic herring, adult fish from the Western Baltic spring spawning (WBSS) population perform feeding migrations into the Skagerrak, and make up a majority of the herring biomass in the Skagerrak in summer (Ruzzante *et al*., 2006; Berg *et al*., 2017). There is, thus, high functional connectivity between three herring populations mechanically admixed in the Skagerrak: Norwegian spring spawners (NSS), North Sea autumn spawners (NSAS), and WBSS (Berg *et al*., 2017). A similar pattern is present in harbour porpoise, where individuals from the western Baltic population appear to migrate into the Kattegat and Skagerrak during the winter (Börjesson *et al*., 1997). Atlantic bluefin tuna (*Thunnus thynnus*) from both the western and eastern stocks also perform feeding migrations into the Skagerrak in late summer-autumn (Aarestrup *et al*., 2022). Active dispersal between seas also appears to be high in European plaice (*Pleuronectes platessa*), where 15- 20% of spawning fish tagged in the North Sea migrate into the Skagerrak, and *vice versa* (Ulrich *et al*., 2017). Similarly, sea trout (*Salmo trutta*) kelts from multiple sites in the North Sea, Skagerrak, and Kattegat appear to migrate long distances, between all three areas, during their time at sea (Kristensen *et al*., 2019).

##### Species with passive dispersal

For European shore crabs (*Carcinus maenas*), there is a clear correlation between larval migratory behaviour and the prevalent tidal regime of each area. When assuming that crab larvae have a circadian vertical migration behaviour, crab larvae passively disperse between the Skagerrak and Kattegat in both directions, and from both seas into the North Sea, while when considering tidal migration behaviour with tidal streams, a dispersal barrier from the North Sea into the Skagerrak seems to form (Jahnke *et al*., 2022). In northern shrimp (*Pandalus borealis*), dispersal of larvae can occur from the North Sea into the Skagerrak (Jorde *et al*., 2015), however, dispersal in the other direction has not been assessed.

For many other species with little to no active migration, dispersal tends to be highly asymmetric in the Skagerrak. In the seaweed tangle (*Laminaria hyperborea*), biophysical modelling shows that spores are mainly transported westward along the Norwegian coast out into the North Sea (Ribeiro *et al*., 2023). For blue mussels (*Mytilus spp.*), larval connectivity within one generation is relatively high from the Kattegat into the Skagerrak, while being lower in the other direction, and practically zero with the Baltic Sea in both directions (Larsson *et al*., 2017; Stuckas *et al*., 2017). A similar pattern, but with lower overall connectivity, is seen in the isopod *Idotea balthica* (De Wit *et al*., 2020).

Connectivity between the Kattegat and Skagerrak is, similarly, high in the northward direction, but practically zero in the southward direction for eelgrass and the phytoplankton *Skeletonema marinoi* (Godhe *et al*., 2013; Jahnke *et al*., 2018).

##### Genetic connectivity

Contemporary genetic connectivity between the Skagerrak and adjacent seas has been inferred indirectly based on estimating rates of gene flow between areas for corkwing wrasse (*Symphodus melops*), star tunicate (*Bothryllus schlosseri*), Pacific oyster (*Crassostrea gigas*), and harbour seal. In all four species, there appears to be some levels of gene flow between the North Sea and Skagerrak (Börjesson *et al*., 1997; Reem *et al*., 2003; Faust *et al*., 2017; Mattingsdal *et al*., 2020). In harbour seals, some levels of gene flow have been inferred between all areas, with gene flow roughly three times higher between the Skagerrak and Kattegat compared to between the Skagerrak and North Sea or Baltic Sea (Goodman *et al*., 1998).

#### 3.2.2. Connectivity within the Skagerrak

Studies focusing on contemporary connectivity on smaller scales, within the Skagerrak, are limited to 28 studies on 10 species – Atlantic cod, Atlantic herring, sea trout, turbot (*Scophthalmus maximus*), broadnosed pipefish (*Syngnathus typhle*), brown crab (*Cancer pagurus*), European lobster (*Homarus gammarus*), blue mussel, harbour seal, and eelgrass. It is worth noting that 12 of the studies were on Atlantic cod, and the remaining 16 studies were distributed across the other nine species.

##### Species with active dispersal

Most studies on mobile species have assessed active dispersal, using mark-recapture or acoustic telemetry. For the two most-studied species, cod and herring, mechanical mixing of genetically differentiated populations is well-documented. These partially sympatric populations or ecotypes have been discovered through population genetic investigations and individual-based clustering approaches, able to unravel complexities and explain some of the statistical noise in tagging/telemetry data.

Most cod tagged in Skagerrak fjords show a tendency toward residency in the fjords (on the timescales studied), but cod of the “offshore ecotype” also show more migratory behaviour out of the fjords (e.g., Pihl & Ulmestrand, 1993; Knutsen *et al*., 2011; Barth *et al*., 2019). Dispersal distances of cod, thus, range from a few to hundreds of kilometres (Svedäng *et al*., 2007). Population genetic studies further support that functional connectivity is high between the offshore and coastal/fjord ecotypes, as they are sympatric on small spatial scales in the Skagerrak (Knutsen *et al*., 2018; Henriksson *et al*., 2023) and even spawn in the same areas, to some extent (Jorde *et al*., 2018; Svedäng *et al*., 2019).

Studies on herring in the artificial estuary Landvikvannet in southern Norway show that three distinct populations coexist in the area: a “local” population (potentially originating from the western Baltic spring spawner population), and small proportions of oceanic Norwegian spring spawners and coastal Skagerrak spring spawners entering the lake in March-April (Eggers *et al*., 2014; 2015). Interestingly, even the “local” herring do not appear to be resident in this “fjord”, but instead migrate to other coastal areas after the spawning season (Eggers *et al*., 2015).

Similar to cod, sea trout in a Norwegian fjord appear to consist of two behavioural phenotypes coexisting in the same habitat – resident “stayers”, and “dispersers” migrating out of the fjord (Thorbjørnsen *et al*., 2019). It is unclear, however, if these constitute different populations (Thorbjørnsen *et al*., 2019). Two migratory phenotypes are also present in brown crab, but here it is the sexes that differ. Males are more-or-less stationary, while females tend to migrate longer distances, sometimes even offshore (Karlsson *et al*., 1996; Ungfors *et al*., 2007).

In contrast, European lobsters, regardless of sex, are very site attached, with adult movement generally limited to less than 1 km (Moland *et al*., 2011a; 2011b; Wiig *et al*., 2013), although some recaptures have also been made up to 25 kilometres away from the release sites (Thorbjørnsen *et al*., 2018).

Juvenile turbot are also relatively stationary along the Norwegian Skagerrak coast, generally recaptured at coastal sites less than 50 kilometres away from the release sites (Bergstad *et al*., 1997). Harbour seals, similarly, had high site fidelity, with individuals dispersing less than 32 km away from their birthplace off the Swedish Skagerrak coast (Härkönen & Hårding, 2001). The mobile species with the lowest dispersal potential out of the species studied, broadnosed pipefish, has a spatial decorrelation scale of 2.4 kilometres, indicating demographic connectivity along the coast is very low (Knutsen *et al*., 2022).

##### Species with passive dispersal

For passively dispersing species or life stages, local retention of eggs and propagules generated inside Skagerrak fjords is common. This is the case for Atlantic cod (Espeland *et al*., 2007; Cianelli *et al*., 2010; Knutsen *et al*., 2011), blue mussel (Pastor *et al*., 2021), and, to some extent, eelgrass (Jahnke *et al*., 2020). For the two assessed species with sessile adult life stages, blue mussel and eelgrass, passive dispersal of propagules along the coast appears to be within 10-45 kilometres (Stuckas *et al*., 2017; Jahnke *et al*., 2020). Interestingly, blue mussel larvae do not seem to passively disperse between the western and eastern Skagerrak or Kattegat (Stuckas *et al*., 2017).

### 3.3. Geographic connectivity barriers

Out of the 12 publications assessing connectivity barriers, two publications found no barrier in the Skagerrak or to the adjacent North Sea and Kattegat, for Pacific oyster (Faust *et al*., 2017) and goldsinny wrasse (*Ctenolabrus rupestris*; Jansson *et al*., 2017). Seven publications describe a barrier located between the North Sea and the Skagerrak, for ballan wrasse (*Labrus bergylta*; Seljestad *et al*., 2020), corkwing wrasse (Blanco Gonzalez *et al*., 2016; Faust *et al*., 2018; 2021; Mattingsdal *et al*., 2020), broadnosed pipefish (Knutsen *et al*., 2022), and shore crab (Moksnes *et al*., 2014). One publication found a barrier for eelgrass between the Kattegat and the Skagerrak (Jahnke *et al*., 2018). The only geographic barriers identified within the Skagerrak were within fjord systems for the passively dispersing eelgrass (Jahnke *et al*., 2020) and blue mussel (Pastor *et al*., 2021).

The re-analysis of genetic distance estimates with BARRIER inferred three main categories of genetic connectivity barriers within the study area. Similar to previous studies, we inferred genetic barriers on the southwestern tip of Norway, between the Skagerrak and the North Sea (Figure 5A). The barrier is shared among ballan wrasse, corkwing wrasse (based on four different studies), pipefish, and tangle. Barriers in this area were also indicated in several other species (Atlantic herring, goldsinny wrasse, lumpfish (*Cyclopterus lumpus*), sea trout, sprat (*Sprattus sprattus*), Norway lobster (*Nephrops norvegicus*), and northern shrimp), but with lower support (Figure 5B-C; thin lines). Other species (Atlantic cod, sand goby (*Pomatoschistus gobius*), sea trout, sprat, European lobster, Norway lobster, northern shrimp, and eelgrass) show various forms of barriers between fjords or coastal locations (Figure 5B). Lastly, many species (Atlantic herring, goldsinny wrasse, lumpfish, cod, plaice, sea trout, and common sole [*Solea solea*]) also have inferred genetic barriers in the southern parts of the area, both between the Skagerrak and Kattegat, and within the Kattegat (Figure 5C). Most of the genetic barriers in the south are more-or-less latitudinal, separating northern and southern populations. Sea trout is an interesting exception to this, as the strongest barrier is between western and eastern populations in the Skagerrak-Kattegat.

**Figure 5.**
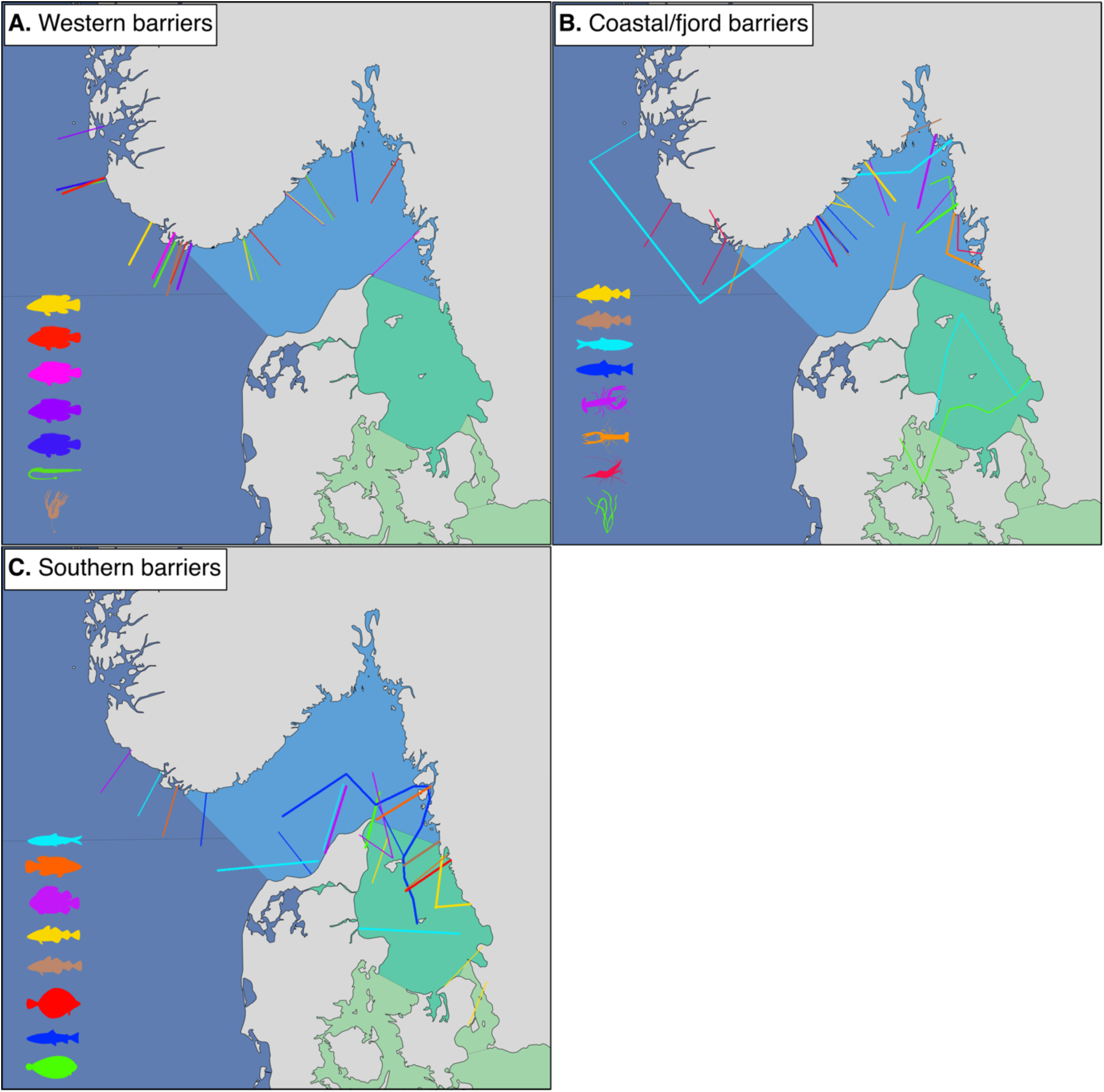
Inferred genetic barriers from the re-analysis of genetic divergence estimates in BARRIER, displayed as lines on maps. The line thickness indicates the level of support for each barrier, and different colours indicate different species and studies. The subplots group studies based on the location of the main inferred genetic barrier: **A)** off southwest Norway, **B)** between coastal locations or fjords, and **C)** in the southern Skagerrak/Kattegat. A key for the species symbols is found in Figure 2E.

### 3.4. Population structure

#### 3.4.1. Population structure against adjacent seas

Based on the reviewed scientific literature, divergence between Skagerrak populations and adjacent populations in the North Sea, Kattegat, and Baltic Sea seems to be the rule rather than the exception (Figure 6). Divergence between the North Sea and Skagerrak has been assessed in the largest number of species (n=38), and the populations diverge in 68 % of these species. The corresponding numbers for the Kattegat (n=26) and Baltic (n=18) are 62 and 89 %, respectively. Note that divergence has not been assessed between all areas in all species, either because of a lack of studies or because the species is not present in all areas.

**Figure 6.**
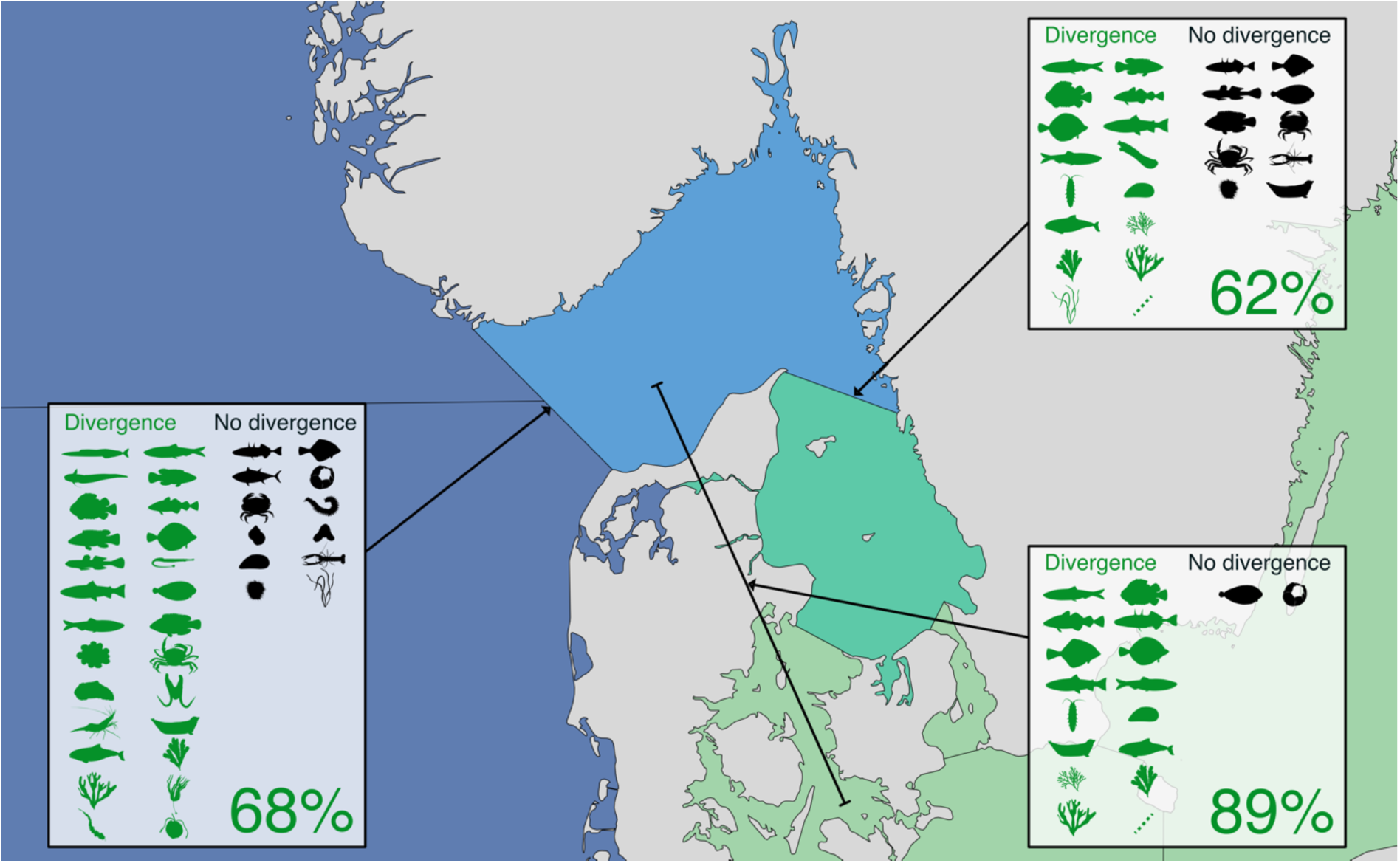
Population structure between the Skagerrak and the adjacent North Sea, Kattegat, and Baltic Sea. The boxes show the species for which population structure between two areas has been assessed, and the proportion of species for which population structure has been found (in green). A key for the species symbols is found in Figure 2E.

In some species, the Skagerrak populations are divergent from populations in all three adjacent areas – Atlantic herring (e.g., Ruzzante *et al*., 2006; Atmore *et al*., 2022; Bekkevold *et al*., 2023), lumpfish (Jansson *et al*., 2023), Atlantic cod (Barth *et al*., 2019), European plaice (Ulrich *et al*., 2017), sea trout (Bekkevold *et al*., 2020), harbour porpoise (Lah *et al*., 2016), toothed wrack (*Fucus serratus*; Coyer *et al*., 2003), and bladderwrack (*Fucus vesiculosus*; Pereyra *et al*., 2023).

In many other species, one or more areas have not been compared to the Skagerrak, however, differences have been found between all of the areas that have been compared. These species include sandeel (*Ammodytes marinus*; Wright *et al*., 2018a), roundnose grenadier (*Coryphaenoides rupestris*; DeLaval *et al*., 2018), goldsinny wrasse (Jansson *et al*., 2023), ballan wrasse (Seljestad *et al*., 2020), broadnosed pipefish (Knutsen *et al*., 2022), star tunicate (Reem *et al*., 2013), Pacific oyster (d’Auriac *et al*., 2017; Faust *et al*., 2017), northern shrimp (Knutsen *et al*., 2015), vase tunicate (*Ciona intestinalis*; Johannesson *et al*., 2018), the flatworm *Gyrodactylus gondae* (Huyse *et al*., 2017), *Idotea balthica* (De Wit *et al*., 2020), tangle (*Laminaria hyperborea*; Evankow *et al*., 2019), sugar kelp (*Saccharina latissima*; Evankow *et al*., 2019; Ribeiro *et al*., 2022), the red alga *Ceramium tenuicorne* (Gabrielsen *et al*., 2002), and the phytoplankton *Chrysocromulina polylepis* (Edvardsen *et al*., 1998) and *Skeletonema marinoi* (Godhe *et al*., 2010; 2013).

For yet other species, there is no known population divergence between any of the areas, e.g., bay barnacle (*Balanus improvisus*; Wrange *et al*., 2016), brown crab (Ungfors *et al*., 2009), Norway lobster (Westgaard *et al*., 2023), rough periwinkle (*Littorina saxatilis*; Panova et a., 2011), the annelid *Hesionides arenaria* (Schmidt *et al*., 2000), warty comb jelly (*Mnemiopsis leidyi*; Verwimp *et al*., 2020), and green sea urchin (*Strongylocentrotus droebachiensis*; Norderhaug *et al*., 2016).

#### 3.4.2. Population structure within the Skagerrak

Population structure within the Skagerrak is slightly less common than divergence from other adjacent areas. However, population structure within the Skagerrak appears to be the rule rather than the exception, with 57 % of the assessed species exhibiting population structure in this region (Figure 7A).

**Figure 7.**
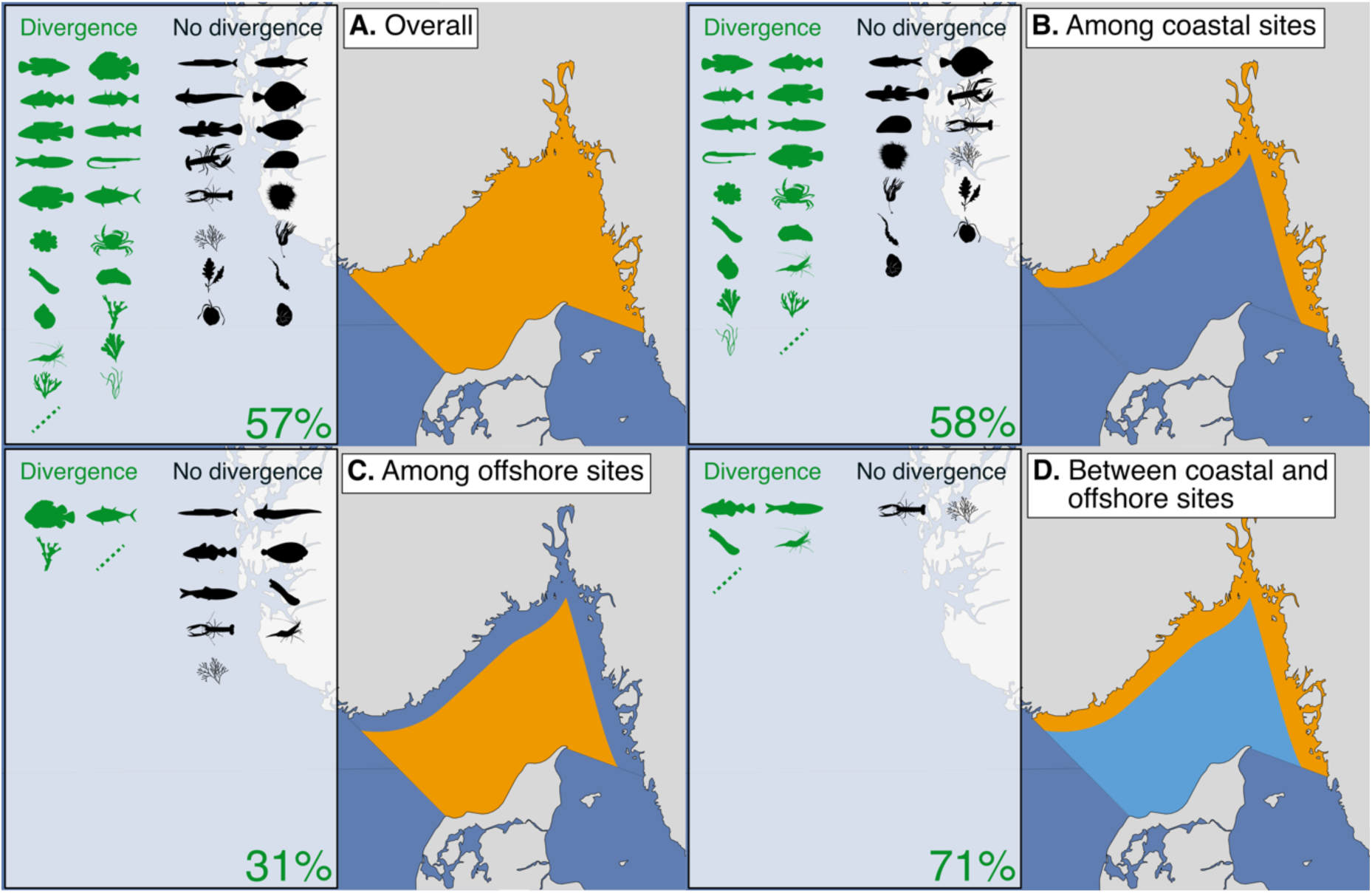
Consensus population structure within the Skagerrak for the species assessed in the scientific literature. Subplots show whether any population structure has been found **A)** broadly within the Skagerrak, **B)** among coastal sites, **C)** among offshore sites, or **D)** between coastal and offshore sites. The boxes show which species have been assessed, and the proportion of these species for which population structure has been found (in green). A key for the species symbols is found in Figure 2E.

The most studied form of divergence is among coastal populations, which is found in 58 % of the assessed species (Figure 7B). These include all three labrid species (Blanco Gonzalez *et al*., 2016; Seljestad *et al*., 2020; Jansson *et al*., 2023), Atlantic cod (e.g., Sodeland *et al*., 2022), three-spined stickleback (*Gasterosteus aculeatus*; De Faveri *et al*., 2013), sea trout (Bekkevold *et al*., 2020), sprat (Quintela *et al*., 2020; 2021; Saltalamacchia *et al*., 2022), broadnosed pipefish (Knutsen *et al*., 2022), star tunicate (Reem *et al*., 2013), shore crab (Jahnke *et al*., 2022), vase tunicate (Johannesson *et al*., 2019), pacific oyster (d’Auriac *et al*., 2017; Faust *et al*., 2017), rough periwinkle (Mäkinen *et al*., 2008; Ravinet *et al*., 2017), northern shrimp (Knutsen *et al*., 2015), toothed wrack (Coyer *et al*., 2003), bladderwrack (Tatarenkov *et al*., 2007; Pereyra *et al*., 2023), eelgrass (Jahnke *et al*., 2018; 2020), and *Skeletonema marinoi* (Godhe *et al*., 2013).

Divergence among offshore populations is rarer (31 %), and only described in four of the thirteen assessed species (Figure 7C): lumpfish (Jansson *et al*., 2023), Atlantic Bluefin tuna (Aarestrup *et al*., 2022), the cold-water coral *Lophelia pertusa* (Dahl *et al*., 2012), and the phytoplankton *Skeletonema marinoi* (Godhe *et al*., 2010; 2013).

An apparently more common form of divergence is between coastal and offshore populations, which is found in five (71 %) out of seven assessed species (Figure 7D). It has been described for Atlantic cod (e.g., Knutsen *et al*., 2018; Henriksson *et al*., 2023), sprat (e.g., Quintela *et al*., 2021; Saltalamacchia *et al*., 2022), vase tunicate (Johannesson *et al*., 2019), northern shrimp (Knutsen *et al*., 2015), and *Skeletonema marinoi* (Godhe *et al*., 2010; 2013). Despite being commonly found, this form of divergence has been studied in the fewest species (n=7).

The species with no clear evidence for population structure within the Skagerrak are: Atlantic herring (e.g., Han *et al*., 2020; Berg *et al*., 2022), roundnose grenadier (Delaval *et al*., 2018), European plaice (Ulrich *et al*., 2017), sand goby (Leder *et al*., 2021), common sole (Cuveliers *et al*., 2012), European lobster (Huserbråten *et al*., 2013), blue mussel (Larsson *et al*., 2017; Pastor *et al*., 2021), Norway lobster (Westgaard *et al*., 2023), green sea urchin (Norderhaug *et al*., 2016), *Ceramium tenuicorne* (Gabrielson *et al*., 2002), tangle (Evankow *et al*., 2019), *Phycodrys rubens* (Van Oppen *et al*., 1995), sugar kelp (e.g., Ribeiro *et al*., 2022), *Chrysochromulina polylepis* (Edvardsen *et al*., 1998), and *Nonionella stella* (Deldicq *et al*., 2019).

## 4. Discussion

### 4.1. Current state of knowledge

Our review shows that the functional connectivity of marine populations in the Skagerrak tends to be high in most species, both within the Skagerrak and with adjacent seas. Meanwhile, population structure is evident in a majority of Skagerrak species. Most species have populations in the Skagerrak that are genetically and/or morphologically distinct from those in adjacent seas, and several species also display finer-scale structure with multiple distinct populations within Skagerrak. Most population structure within the Skagerrak tends to be associated with the coastal habitat, and fjords in particular. Divergence is common among coastal populations, but rarer among offshore populations. In line with the population genetic findings, small-scale connectivity studies indicate local retention of propagules in several fjords, and high prevalence of resident behaviour in multiple mobile species. This stands in contrast to the high functional connectivity inferred on the large scale, between the Skagerrak and the North Sea, Kattegat, and Baltic Sea. These findings imply that management of marine biodiversity in this region needs to be considered on the scale of Skagerrak or finer.

The somewhat counterintuitive combination of both high connectivity and evident population structure illustrates the importance of appreciating that there are different types of connectivity of relevance in shaping population structure in marine species. Population genetic theory tells us that very low numbers of migrants are sufficient to erode most genetic population structure (Slatkin, 1987). The maintenance of genetically distinct populations within species, despite high functional connectivity, implies that factors other than contemporary dispersal and migration may limit the levels of gene flow. Populations may share feeding grounds or nursery areas during parts of the year or across certain life stages, while behavioural differences may ensure that spawning occurs in allopatry or at different times, as is the case in Atlantic herring (Ruzzante *et al*., 2006) and cod (André *et al*., 2016). Genetic differences, themselves, may also put limitations on the extent of gene flow across populations, for instance if hybrid offspring are inviable (Irwin, 2020). Differences in the genetic architecture of populations may lead to heterogeneity in the levels of gene-flow in different parts of the genome (Harrison & Larson, 2016). For instance, chromosomal inversions, which have been described for Atlantic herring and cod in the Skagerrak, may enable sympatric genetic divergence despite ongoing gene-flow (Han *et al*., 2020; Sodeland *et al*., 2022). The current transition into population genomic analysis using SNP loci may provide more insight into the importance of both genetic architecture and adaptive genetic divergence in limiting hybridisation in marine species.

Importantly, though, the high functional connectivity between populations, itself, also has relevance for conservation management. The partial sympatry of genetically differentiated populations means a mechanical mix of individuals from different populations can coexist in the same place during different parts of the year. This fact complicates the use of spatial methods to delineate populations or define the borders of MPAs.

### 4.2. Knowledge gaps

Our review reveals that studies on demographic connectivity in the Skagerrak are relatively rare. This is a notable finding, since it may be the most important form of connectivity for spatial MPA management. It is of critical importance to not only know how individuals move between areas (functional connectivity), but to also relate this movement to population growth and source-sink dynamics (demographic connectivity). A key challenge in this regard is to accurately and consistently define when an individual has fully recruited into the new population, in demographic terms. Another issue is that the effects of immigration and emigration depends on the population size of the receiving populations. Accurate estimation of demographic connectivity requires knowledge on both migration rates and intrinsic demographic rates of the populations (Lowe & Allendorf, 2010). Such estimations fundamentally require accurate identification and definition of population A and B, which can be challenging if knowledge on the underlying population structure is lacking. Even in species with accurately defined population structure, estimates of population sizes may be unavailable. This illustrates how the concepts of population structure and connectivity are deeply interconnected also for management applications. However, despite their importance, this type of information is generally lacking within marine fisheries and biodiversity management.

We found that there is an increasing number of studies that jointly assess both genetic and functional connectivity in the Skagerrak. This is a promising way forward from considering population genetics separately from tagging or biophysical modelling studies – the former focusing solely on identifying population structure or adaptive genetic variance within species, and the latter two focusing solely on dispersal behaviour. Studying both in concert enables connecting genetic origins to phenotypes, hence, moving past merely identifying populations, and towards understanding the ecological relevance of population structure.

There is a tendency that species with a larger number of studies are more likely to have population structure described for them. There may be several underlying causes for this pattern. On the one hand, population structure is more likely to be discovered the more studies have been performed on the species. On the other hand, species for which population structure has already been described may be a more attractive target for further studies. This tendency can lead to knowledge being restricted to a few model species, with limited possibilities for broader comparisons of patterns across taxa and applicability of concepts to other systems. Currently, there is an obvious taxonomic bias toward fish species within connectivity research in the Skagerrak (Figure 2B). The large amount of studies on model species such as Atlantic cod has provided key insights into marine evolutionary biology and connectivity research, which have helped move the entire research field forward. For instance, within Atlantic cod, there has been a gradual transition from previously estimating genetic divergence at the sample-level to now using individual-based clustering methods to infer genetic origins. This has led to the discovery that individuals of multiple genetic origins may frequent the same geographic area. This may be the case in other species as well, but further study is then needed. At present, however, knowledge on 48 % of the species covered in this review is limited to single studies (22 out of 46 species), often confined to small geographic areas. In fact, information about population structure and connectivity is only available for a small fraction of the thousands of species documented in the Skagerrak. Thus, expanding the knowledge on population structure and connectivity to other, non-fish taxa is of great importance. This is especially important to enable generalising patterns of intraspecific divergence and connectivity across taxa, to avoid leaving the management of these non-model species to guesswork. Furthermore, these understudied taxa may perform important ecosystem functions, such as being ecosystem engineers or forming the base of the food chain.

In species with low numbers of publications, the knowledge on population structure and connectivity may also be restricted to a smaller geographic area. The reduced geographic representation of certain species limits the ability to find population structure and connectivity barriers. Also, the patterns that have been described in the species may not be the biologically most relevant, but rather artifacts from the geographic sampling design. For instance, in our BARRIER analysis, different studies on the same species inferred different genetic barriers, most likely due to differences in the geographic region sampled (see, e.g., sea trout and Atlantic cod in Figures 5B-C). Limited geographic extent may also bias analyses of connectivity. For example, the dispersal distances reported in small-scale tagging studies may depend on the extent of the telemetry arrays or the regions of recaptures. In some studies, individuals dispersing outside of the telemetry array are completely excluded from further analysis.

This may be justifiable based on the scope of the original study, but it likely underestimates dispersal distances.

### 4.3. Implications for management

By reviewing the available scientific literature, we show that a majority of species have populations inhabiting the Skagerrak that are genetically and/or morphologically distinct from surrounding populations in the North Sea, Kattegat, and Baltic Sea. Additionally, many species also have several distinct populations within the Skagerrak. Despite this, functional connectivity on the large scale is high in most species, meaning individuals from several populations may coexist in certain areas during parts of the year, especially in highly mobile taxa such as the more mobile fish species. Working according to this connectivity “rule book” is likely essential to achieving sustainable management of intraspecific biodiversity in the Skagerrak. However, with these findings come considerable challenges.

The high contemporary connectivity with adjacent seas on the large scale supports the notion of the Skagerrak as a transition zone between the North Sea and the Baltic Sea. Multiple species have populations dispersing in and out of the Skagerrak at different times. Management of marine populations in the area, thus, cannot view the Skagerrak as an entirely isolated system, but needs to take large-scale connectivity into consideration. As shown herein, however, the Skagerrak itself is not homogeneous. Most species have multiple differentiated populations, particularly along the coast, which are unique to the Skagerrak. Management of such species should be on a much finer geographic scale than the entire Skagerrak to preserve unique populations – often on the scale of 10s of kilometres, or of individual fjords. The Skagerrak is, thus, more than just a transition zone – it is also a unique marginal sea requiring special attention from management, to preserve both coastal and offshore populations.

For many of these species, particularly in more vagile fish species, there is also temporal variability in population assemblages. The sympatry of multiple populations within a species in a given area poses a significant challenge for spatial methods used to delineate management units. Management of these species should establish practices suitable for mixed-stock management. For instance, in mixed-stock fisheries, the relative proportions in catches over time should be monitored, for instance using population genetic tools, with management decisions taken expediently according to the relative population sizes. Management, thus, needs to be both temporal and spatial.

Adding to this point, the fields of connectivity and population structure are growing, gaining more research attention and more utility in legislation. With this development, we are gaining increasingly detailed knowledge, as well as improved taxonomic and geographic representation at both large and small scales. Consequently, management strategies need to be both spatiotemporally sensitive, and flexible enough to adapt to new scientific findings. For instance, management programs for monitoring genetic diversity (e.g., Mastretta-Yanes *et al*., 2024), and real-time genetic monitoring of fisheries catches (e.g., Dahle *et al*., 2018) have been suggested to improve management practices, by enabling agile management in response to updated information on intraspecific diversity and connectivity. Incorporating these monitoring tools in management would also aid in the estimation of population sizes, fundamental to analyses of demographic connectivity, which are lacking in this region.

Ideally, if knowledge is lacking for a species of interest, knowledge on a species with similar life-history may be used as guide or proxy. However, we have found that different taxa are so unequally represented that patterns cannot justifiably be generalised across species. While the knowledge on some species may suffice for informing the design of spatial management measures, improvement can be gained from temporal and spatial connectivity assessment on a per species basis. Synthesising knowledge on population structure and connectivity for multiple species, as we have done here, helps paint the broader picture, but new and more detailed knowledge is generated for each species continuously. Hence, adaptive management approaches combining spatial and temporal management are more likely to succeed in establishing a robust and future-proof management regime for biodiversity in the Skagerrak.

Below we briefly summarise our main recommendations for management:

- Management of marine biodiversity in the Skagerrak needs to be based on knowledge about species population structure and connectivity.
- Management should be fine-scaled enough to capture population structure within the Skagerrak, often on the scale of 10s of km, especially along the coast and within fjords.
- Fisheries management, MPA design and marine spatial planning need to consider both coastal and offshore marine areas.
- Management needs to consider that different populations may coexist at certain times in a given area. This is especially relevant in fisheries management, when different stocks coexist, and where genetic mixed-stock analysis should be implemented to disentangle and estimate the proportions of the different stocks.
- More information on population structure and connectivity is needed, both for sessile and mobile species.
- Adaptive strategies that incorporate both spatial and temporal management are more likely to succeed in creating a robust and future-proof biodiversity management in the Skagerrak.

## 5. Conclusions

The finding of differentiated populations despite high functional connectivity, although counterintuitive, is not specific to the Skagerrak. Differentiated but sympatric populations are found in marine species in many other regions around the world, running the same risks of mismanagement unless accounted for (e.g., Le Moan *et al*., 2016; Moore *et al*., 2021; Diaz-Arce *et al*., 2024).

Therefore, we believe that the patterns inferred here, as well as our recommendations for management, are likely to be more broadly applicable also to other marine systems. In this review, we have described overarching patterns of connectivity and population structure in a marginal sea, enabling marine wildlife management to better account for, conserve, and restore biodiversity – on all levels.

## Author contributions

Conceptualisation: SH, CA, MJ, PDW, EM. Investigation: SH, CA, CB, PDW, MJ, PEJ, GS. Formal analysis: SH, PEJ. Visualisation: SH. Writing – Original Draft: SH. Writing – Review and Editing: CA, CB, PDW, MJ, PEJ, HK, EM, GS. Funding acquisition: CA, EM.

