## Supplementary Tables S1-2 for "Connectivity and population structure in a marginal sea – a review"

**Table S1.** Species and publications included in the BARRIER analysis. The number of barriers was defined *a priori* based on the number of sampling localities in the Skagerrak, according to Table S2.

| **Species** | **Localities** | **Barriers** | **Source:** |
|---|---|---|---|
| ballanwrasse | 7 | 3 | Seljestad *et al.* 2020 |
| codA | 5 | 3 | Jorde *et al.* 2007 |
| codB | 6 | 3 | Knutsen *et al.* 2003 |
| codC | 7 | 3 | André *et al.* 2016 |
| codD | 6 (3) | 1 | Svedäng *et al.* 2019 |
| corkwingA | 11 | 4 | Blanco_Gonzalez *et al.* 2016 |
| corkwingB | 4 | 2 | Faust *et al.* 2018 |
| corkwingC | 6 | 3 | Faust *et al.* 2021 |
| corkwingD | 5 | 3 | Mattingsdal *et al.* 2020 |
| goldsinny wrasse | 9 | 3 | Jansson *et al.* 2023 |
| herring | 6 | 3 | Bekkevold *et al.* 2015 |
| laminaria hyperborea | 4 | 2 | Evankow *et al.* 2019 |
| lobster | 8 | 3 | Huserbråten *et al.* 2013 |
| lumpfish | 10 | 3 | Jansson *et al.* 2023 |
| norwaylobster | 8 | 3 | Westgaard *et al.* 2023 |
| pipefish | 14 | 4 | Knutsen *et al.* 2022 |
| plaice | 3 | 1 | Ulrich *et al.* 2017 |
| seagrass | 22 | 5 | Jahnke *et al.* 2018 |
| seatroutA | 7 | 3 | Knutsen *et al.* 2001 |
| seatroutB | 42 | 6 | Bekkevold *et al.* 2020 |
| shrimp | 18 | 4 | Knutsen *et al.* 2015 |
| sole | 3 | 1 | Cuveliers *et al.* 2012 |
| sprat | 12 | 4 | Quintela *et al.* 2020 |
| tunicate | 6 | 2 | Johannesson *et al.* 2018 |

**Table S2.** The relationship between the number of sampling localities and the number of barriers defined *a priori* in BARRIER.

| **Number of localities** | **Number of barriers** |
|---|---|
| 3 | 1 |
| 4 | 2 |
| 5-10 | 3 |
| 11-20 | 4 |
| 21-30 | 5 |
| >30 | 6 |
